# Bhutan sequences complete the range-wide mitochondrial phylogeography for red pandas, *Ailurus fulgens*

**DOI:** 10.1101/2025.10.13.682096

**Authors:** Lucy A. Dueck

**Affiliations:** University of Georgia’s Savannah River Ecology Laboratory, Aiken, SC 29802, USA (retired); Athens, TX 75752, USA

**Keywords:** endangered species, gene flow, genetic cline, matriarchal population structure

## Abstract

Red pandas, endangered mammals inhabiting an elongated montane region in south-central Asia, are up for IUCN Red List reassessment in 2025. One issue determining their status is species taxonomy. A recent mtDNA analysis compiled from several previous studies found they were united as one species, with weak support for a western sublineage and four haplogroups correlated with specific geographic regions. However, the crucial section of their midrange, Bhutan, was absent from that analysis. Now, sequences from red pandas in Bhutan have been added to that dataset and reanalyzed here. Phylogenetic analyses of these data, 91 haplotypes from ∼770 samples representing the entire red panda range, indicate again that red pandas are only one species, based on mtDNA. Bhutan hosted three of the previously defined four haplogroups and both putative subspecies (red and pink *A. f. fulgens*, orange *A. f. styani*). However, with the addition of the Bhutan missing link, the haplogroups graded even more into one another phylogeographically across their range, effectively displaying a genetic cline. Additionally, no large geographically specific subclade representing a subspecies was significantly supported within the large clade grouping all red pandas as one species. This study further reinforces the conservation priority of encouraging gene flow via protecting prime habitat areas internationally, connected by wildlife corridors and transboundary cooperation, allowing natural evolution for species survival.

## Introduction

As an elusive, rare, arboreal mammal inhabiting select montane areas over a long, narrow range across several countries in south-central Asia, red pandas (*Ailurus fulgens*, Cuvier 1825) have been difficult to study in the wild. But understanding their ecology, behavior, population structure, genetic diversity, and patterns of gene flow has become imperative to properly protect them from threats to this species’ survival. Scientists have contributed studies on these topics that are specific to individual countries or regions within, but none known have thoroughly accumulated enough knowledge about any one topic that encompasses the entire geographic range of red pandas. And partially due to this segmentation, and to separate descriptions from opposite ends of the range 77 years apart, red pandas have been considered two different subspecies, *A. f. fulgens* and *A. f. styani* (Roberts 1982), or even separate species (Hu et al., 2020).

However, it was possible to combine results of mitochondrial DNA control region (mtDNA-CR) analyses from multiple studies across the range (China and Myanmar - Li et al. 2005, Hu et al. 2011; India - Dalui et al. 2021; Nepal WRPs - Dueck and Steffens 2022) that narrowed the sampling gap to provide a broader perspective on matriarchal diversity, population structure, and gene flow for all red pandas except those from Bhutan (Dueck, 2025). Now those missing data have recently become available through a comprehensive study by the government of Bhutan and carried out by the Zoological Survey of India (Dhendup et al., 2023), combining mtDNA and microsatellite analyses. As a result of this survey, red panda mtDNA sequences were deposited in GenBank and accessible for inclusion with the rest of the known, comparable data used by Dueck (2025).

Therefore, the present study aims to update the mitochondrial phylogeography of all native red pandas by analysis based on complete sampling throughout the entire range with addition of the Bhutan information.

## Methods and Materials

The 56 red panda mtDNA sequences (528 bp) from Bhutan submitted by the Zoological Survey of India (#OR666797.1 – OR666852.1) were accessed in GenBank and appeared to represent 432 samples successfully extracted from a scat survey, according to the report (Dhendup et al., 2023). They were added to the full 75-haplotype dataset of Dueck (2025) representing 340 samples, including from the above studies, zoo SSP (Dueck 2021), and two NCBI reference standards (Arnason et al. 2007, Yonezawa et al. 2007). They were aligned in the ClustalW option of MEGA 4.0.2 (Tamura et al. 2007), then trimmed to a corresponding 340 bp length. The Bhutan sequences were subsequently consolidated to 22 haplotypes by comparing with this dataset in unrooted phylogenetic analyses performed in MEGA 4 using the neighbor-joining method (NJ; Saitou and Nei 1987) with 10,000 bootstrap replicates, pairwise deletion, and the Tamura-Nei model to infer evolutionary distances. Duplicates identified in resultant phylograms were visually confirmed in MEGA’s Data Explorer and/or by a NCBI BLAST search. This produced a combined dataset of 91 red panda haplotypes from 772 samples after all duplicates were consolidated.

Two phylogenetic methods were used to examine evolutionary relationships among haplotypes – NJ which is distance based and provides a good approximation of a minimum evolution tree, and the more stringent maximum-likelihood (ML) which is probability based. Red panda haplotypes only were first analyzed together by the NJ method as above to produce an unrooted radial phylogram, which illustrates genetic distance and provides statistical support. After omitting one Bhutan haplotype (see explanation in Results), they were then combined with 10 outgroups from their nearest relatives, the Musteloidea, to root and balance an NJ phylogeny in the form of a condensed (>50% bootstrap support) circular cladogram.

These remaining 90 red panda haplotypes, representing up to 768 samples, were also analyzed via the ML “slow” method in MEGA 12.0.11 (Kumar et al. 2024), using the HKY model of evolution, 1000 replicates, and one from each of the three musteloid families rooting the analysis to produce a condensed rectangular cladogram. This cladogram was combined with a distribution map of red panda haplotype-groups (haplogroups) as previously defined in Dueck (2025) and determined by associations in the NJ radial phylogram.

## Results

The unrooted, uncondensed, radial NJ phylogram (**Figure 1**), with Bhutan red panda haplotypes denoted by blue dots and subspecies’ reference standards by red and yellow squares (*A. f. fulgens* and *A. f. styani*, respectively), indicates a separation into four previously defined haplogroups (Dueck 2025) – red and pink ovals representing the putative subspecies *A. f. fulgens*, orange and yellow representing *A. f. styani*. Note that all Bhutan haplotypes are broadly distributed across the left half of the tree within orange, red, and pink ovals, thus encompassing both putative subspecies. However, the minimal bootstrap values on the haplogroup branches do <u>not</u> support any division into either haplogroups (5-38%) or subspecies (23%). One haplotype in the pink oval representing four Bhutan sequences (B-7417-7312-7168-7123), as denoted by an asterick**\***, was omitted from further rooted analyses because it contained three abnormal sites identical to most of the musteloids used, which skewed the analyses.

**Figure 1.**
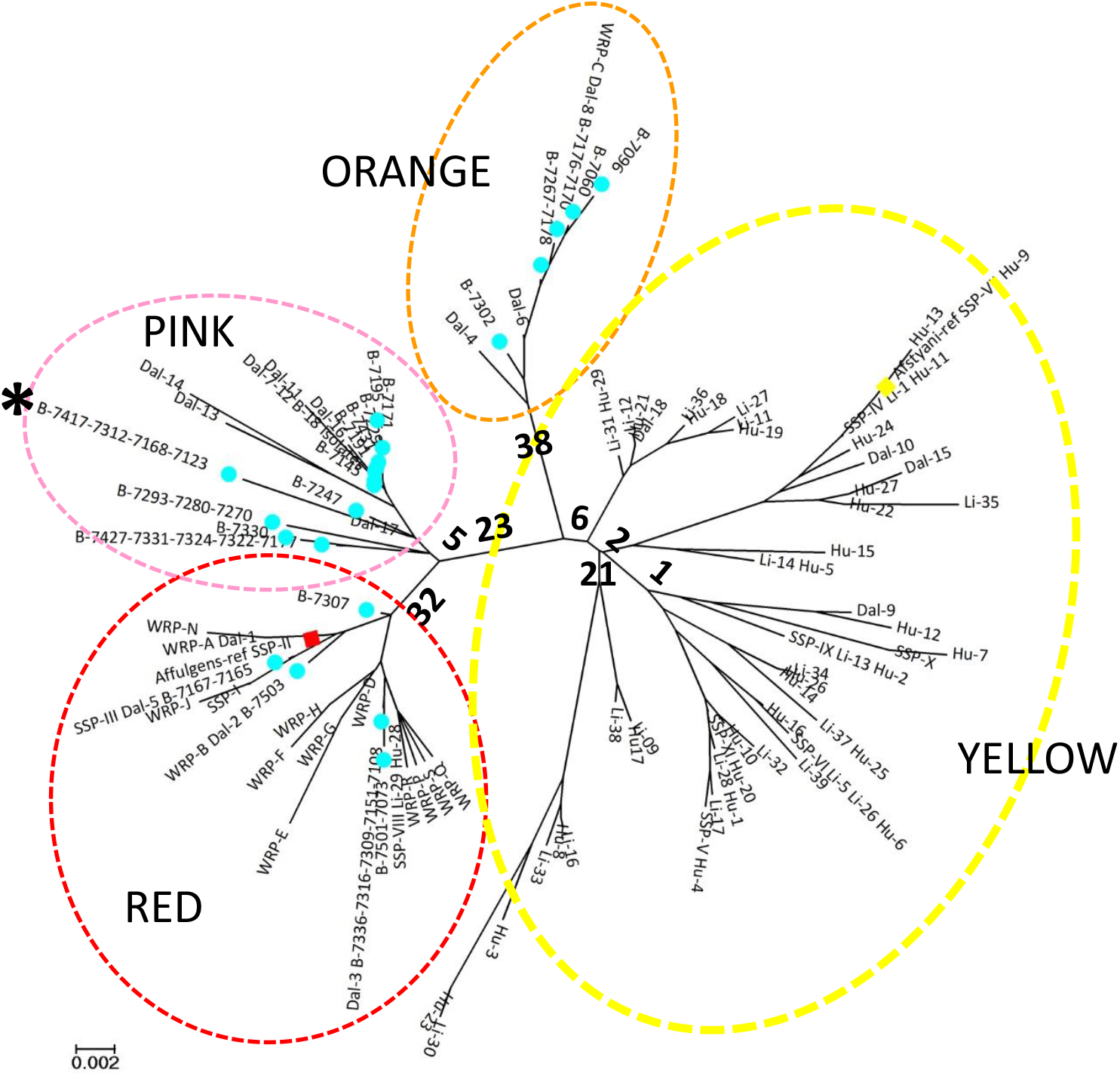
Radial phylogram from unrooted, uncondensed NJ phylogenetic analysis of 91 red panda haplotypes sampled across entire range (*N* = 772), based on 340 bp of mtDNA-CR, computed in MEGA 4. Evolutionary distances computed using Tamura-Nei method, positions containing gaps eliminated only in pairwise comparisons, and 10,000 bootstrap replicates provide support values. The tree is drawn to scale, with branch lengths in the same units as those of the evolutionary distances (base substitutions per site) used to infer the phylogenetic tree. Blue dots indicate haplotypes from Bhutan, red and yellow squares indicate reference standards for putative subspecies. Dashed ovals (red, pink, orange, yellow) indicate haplogroups previously defined in Dueck (2025). Haplotype designated by asterick**\*** was omitted from further rooted analyses.

In the condensed, rooted NJ circular cladogram (**Figure 2**), all 90 red panda haplotypes group together as one species, with 100% bootstrap support. Bhutan and reference standards are denoted by colors as above. There is also no indication of clustering into haplogroups although they are demarcated on this tree as previously defined by the NJ phylogram above. Note that the red haplogroup grades into the pink which grades into the orange then into the yellow haplogroup. There is also no division between putative subspecies.

**Figure 2.**
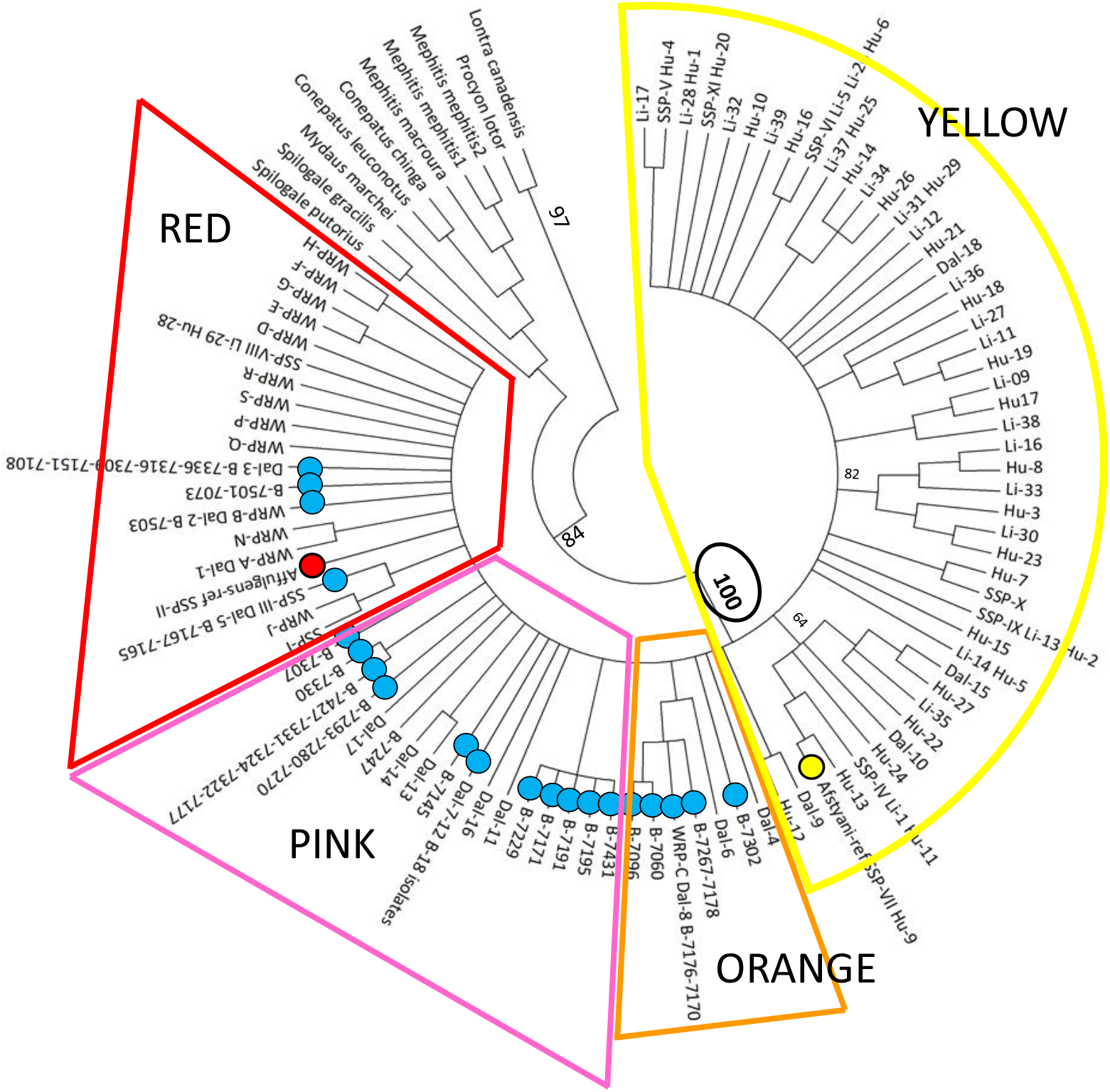
Circular cladogram from condensed NJ phylogenetic analysis of 90 red panda haplotypes sampled across entire range (*N* = ≤768), rooted by 10 outgroups of musteloids, based on 348bp of mtDNA-CR, computed in MEGA 4, using the Tamura-Nei method, pairwise deletion option for gaps, and 10,000 bootstrap replicates to determine support values. Branches with less than 50% support are collapsed. Color designations of dots same as in Figure 1. Although statistically unsupported, the previously defined haplogroups of Dueck (2025) are designated by the same colors and members as determined in Figure 1.

Figure 3 illustrates the relationship between the phylogenetic tree – a condensed, rooted ML rectangular cladogram – and geographic origins of the haplotypes therein. Again, all colors are as denoted above. First, red pandas are fully supported (100%) as a single clade – one species with no subdivision into either haplogroups or subspecies, where these previously defined groupings now grade into one another phylogenetically, based on mtDNA. Second, there is an overall geographic pattern to this distribution which, more or less, correlates with the phylogenetic tree. The (former) red haplogroup was found in western and midrange red pandas from Nepal and Sikkim-W.Bengal (India) - Bhutan, respectively. The (former) pink haplogroup was found in midrange animals from Bhutan and western Arunachal Pradesh (India). Both of these haplogroups were considered belonging to putative *A. f. fulgens*. However, the (former) orange haplogroup was found in the midrange also (Sikkim-W.Bengal and Bhutan), and it clustered previously within *A. f. styani*. All three of these haplogroups were found in Bhutan (although exact sampling locations within are not available), and only west of the Siang River (except one outlier in southeast Tibet). The (former) yellow haplogroup, previously considered *A. f. styani*, was found broadly distributed east of the Siang River in eastern Arunachal Pradesh, Myanmar, and China (Yunnan-Sichan), with one outlier in south-central Tibet. Although some small subclades within the yellow haplogroup have >50% support, no pattern of geographic association was found within them.

**Figure 3.**
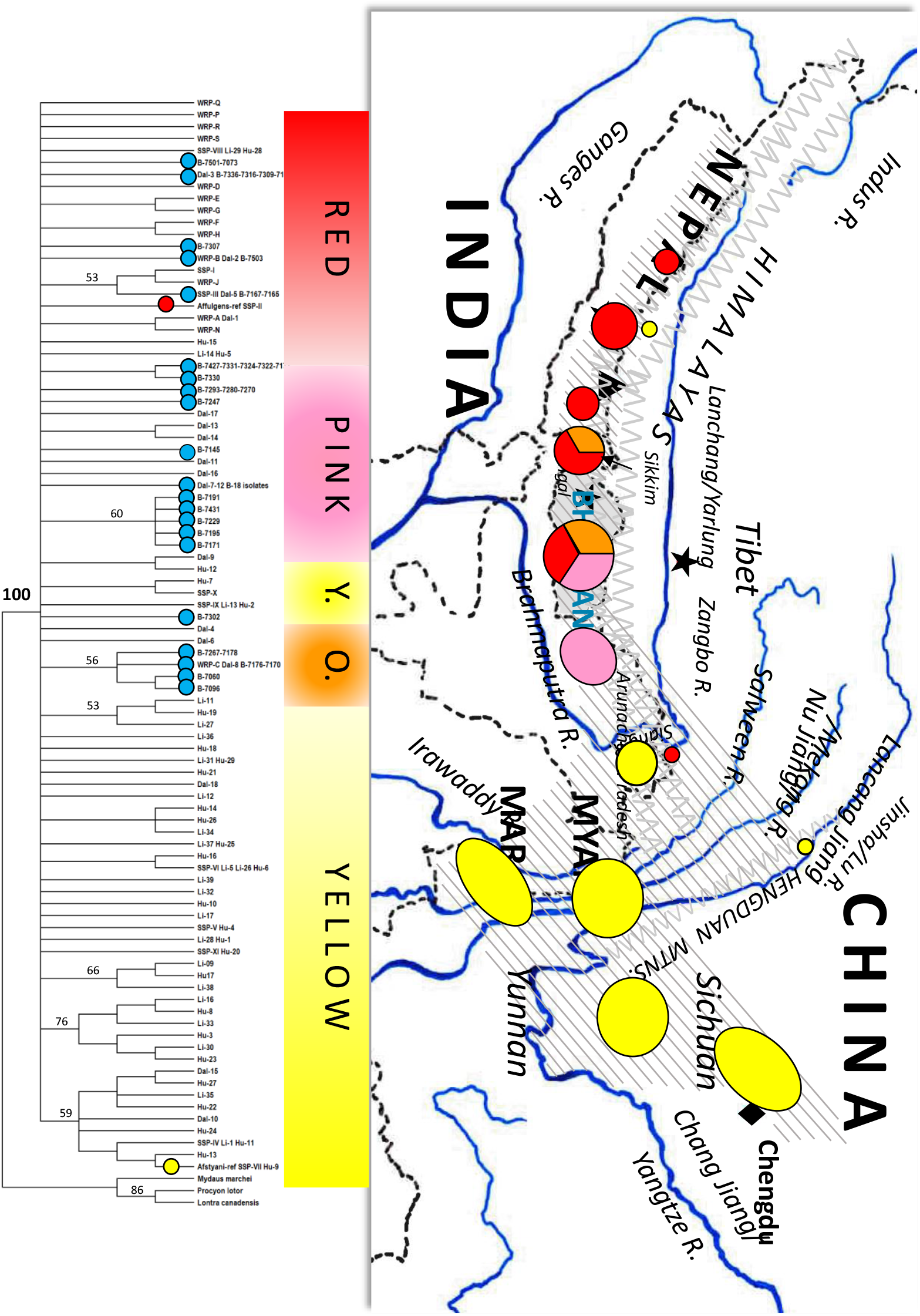
Rectangular cladogram from condensed ML phylogenetic analysis of 90 red panda haplotypes sampled across entire range (*N* = ≤768), rooted by three musteloid outgroups, based on 347bp of mtDNA-CR, computed in MEGA 12 using “slow” method, HKY model of evolution, and 1000 bootstrap replicates for support. Branches with less than 50% support are collapsed. Color designations of dots same as in Figure 1. Although statistically unsupported, the previously defined haplogroups of Dueck (2025) are designated by the same colors and members as determined in Figure 1. The associated map with sampling sites illustrates where the haplogroups from different parts of the phylogenetic tree were obtained. (Note that “pie” pieces from the two divided circles are not proportional to their distribution there.)

## Discussion and Conclusions

The entire range of native red pandas is now genetically represented for the mitochondrial genome by a sample size of more than 750 after combining results of studies conducted from 2005 to 2023. Ninety-one haplotypes were defined from 340 bp of control region sequences. It is likely that these haplotypes could be further divided if longer segments were used, but given this region is hypervariable, it provides a reasonable representation of matriarchal population structure across all the red panda range.

These results indicate that red pandas are genetically only one species, and not significantly subdivided into subspecies or even haplogroups, due to at least female gene flow historically or currently. But there is a gradation of slight differences from one end of their range to the other, implying a genetic cline. Location of the previously-defined *A. f. styani* orange haplogroup in the western midrange and *A. f. fulgens* pink group in the eastern midrange could be the result of early Pleistocene migration, while limitation of the previously-defined *A. f. styani* yellow haplogroup to east of the Siang River could be a function of later glacial timing/location acting as a barrier then. Biogeoclimatic scenarios for this pattern are more thoroughly discussed in Dueck (2025), but further explanation might also be found in behavioral studies. Radiotelemetry work by Bista et al. (2021, 2022) noted that while adult males had 1.5-2 times the territory size of adult females indicating polygyny often linked to male dispersal, two female sub-adults dispersed an average of 21 km while the sub-adult male did not leave, suggesting female-biased natal dispersal. Even if some significant genetic population structure is found among all red pandas from nuclear DNA or Y-chromosome studies, these results of a genetic cline from mtDNA cannot be discounted as part of the whole perspective on their species, as they would indicate a complex system that cannot be simply defined.

As expected, Bhutan acts as a crossroads to host three of four previously-determined haplogroups and both putative subspecies, with the red and pink *A. f. fulgens* found on opposite sides of its borders, and orange *A. f. styani* found on its western border. The country has prioritized respect for nature by establishing many protected areas with good habitat and biological corridors (Dhendup et al. 2023), and thus may act as a refuge for the species. It would also be the ideal location for further studies on the evolutionary basis of this pattern, employing integrative taxonomy. Meanwhile, gene flow should be encouraged throughout the entire range with wildlife corridors, transboundary cooperation, and maintaining red pandas as one diverse species, *Ailurus fulgens*.

## Note

This manuscript was prepared from an addendum to a presentation at the Red Panda Symposium, originally scheduled for November 3-7, 2025, to be considered in the 2025 reassessment of endangered status for red pandas in the IUCN Red List. However, the symposium has been postponed until 2026 (i.e., after that decision will be made), so this information is being provided via written format in October 2025. The completion of mtDNA analyses for all native red pandas would not have been possible without the studies done by the Zoological Survey of India conducted in their midrange.

## Notes

### Competing Interest Statement

The authors have declared no competing interest.

